# Endoplasmic reticulum stress induces a novel Ca^2+^ signalling system initiated by Ca^2+^ microdomains

**DOI:** 10.1101/2020.10.07.328849

**Authors:** Constanza Feliziani, Gonzalo Quasollo, Deborah Holstein, Macarena Fernandez, James C Paton, Adrienne W Paton, James D Lechleiter, Mariana Bollo

**Author notes:** Corresponding author at: Instituto de Investigación Médica M y M Ferreyra, INIMEC-CONICET, Universidad Nacional de Córdoba, Córdoba, Argentina., E-mail address (M. Bollo). Gonzalo Quasollo and Deborah Holstein contributed equally to this study.

## Abstract

The accumulation of unfolded proteins within the Endoplasmic Reticulum (ER) activates a signal transduction pathway termed the <u>u</u>nfolded <u>p</u>rotein response (UPR), which attempts to restore ER homeostasis. If homeostasis cannot be restored, UPR signalling ultimately induces apoptosis. Ca^2+^ depletion in the ER is a potent inducer of ER stress. Despite the ubiquity of Ca^2+^ as intracellular messenger, the precise mechanism (s) by which Ca^2+^ release affects the UPR remains unknown. Use of a genetically encoded Ca^2+^ indicator (GCamP6) that is tethered to the ER membrane, uncovered novel Ca^2+^ signalling events initiated by Ca^2+^ microdomains in human astrocytes under ER stress, as well as in a cell model deficient in all three IP_3_ Receptor isoforms. Pharmacological and molecular studies indicate that these local events are mediated by translocons. Together, these data reveal the existence of a previously unrecognized mechanism by which stressor-mediated Ca^2+^ release regulates ER stress.

## Introduction

An abrupt Ca^2+^ concentration gradient -- over 4 orders of magnitude -- is present in cells between the cytosol and external medium. In the resting state, cytosolic free Ca^2+^ concentration ([Ca^2+^]i) is maintained at 50-100 nM, reflecting a balance between active uptake by Ca^2+^- ATPases and passive release via leak channels. In addition to these fluxes, Ca^2+^-binding capacity of cytosol also plays a role in maintenance of this equilibrium ^1,2^.

Increases in [Ca^2+^]i is a ubiquitous intracellular signal that regulate many cellular processes, including secretion, muscle contraction, neuronal activity, and cell death ^2^. To achieve this versatility, Ca^2+^ signalling is differentiated by temporal, spatial, frequency, and amplitude patterns ^3^. Thus, certain Ca^2+^ events give rise to a highly localised Ca^2+^ increase (microdomains), whereas others generate Ca^2+^ increases that spread through the entire cell (global), often appearing as repetitive waves ^4^. Ca^2+^ microdomains remain localised because Ca^2+^ is buffered before it diffuses into a larger volume ^5,6^, and these local events are generated by Ca^2+^ channel clusters, arranged in discrete membrane domains ^7,8^.

An important spatiotemporal aspect of both global and local Ca^2+^ signalling is the occurrence of a positive feedback process (Ca^2+^-induced Ca^2+^-release; CICR) in intracellular channels -- the basis for their ability to generate repetitive Ca^2+^ spikes and waves ^3^.

The ER is the main Ca^2+^ signalling organelle. In addition to storing and releasing Ca^2+^, the ER plays a critical role in many other processes, such as lipid synthesis, transduction and folding of proteins as well as post-translational protein modification. Alteration of one of these processes can have a strong impact on one of the others. For example, Ca^2+^ depletion in the organelle induces ER stress, resulting in production of misfolded proteins and consequent activation of a protective response termed <u>u</u>nfolded <u>p</u>rotein response (UPR). Inositol-requiring enzyme 1 (IRE1) and protein kinase RNA-like ER kinase (PERK) are transmembrane proteins that reside in the ER and sense stress ^9,10^. The luminal domains of these UPR sensors are in complex with the chaperone protein BiP (<u>b</u>inding immunoglobulin <u>p</u>rotein), a master regulator that maintains them in an inactive state. Under stress conditions, BiP is competitively titrated by the unfolded proteins, leading to oligomerization and UPR sensors activation ^11^. During acute stress responses, PERK and IRE1 activities reduce protein synthesis by, respectively, inhibiting protein transduction and degrading RNAs ^12-14^. Throughout the adaptive phase of the stress response, the ER functions are facilitated by expression of a group of genes including ER chaperones, such as BiP ^15^. If ER stress is persistent and the adaptive mechanisms are not sufficient for restoration of homeostasis, UPR signalling is switched off, inducing cell death by apoptosis ^16^.

We previously demonstrated that calcineurin, a Ca^2+^-dependent protein, promotes cell survival during the acute phase of UPR in human and mouse astrocytes as well as in *Xenopus* oocytes ^17,18^. Expression levels of calcineurin are rapidly increased and it interacts with a cytosolic domain of PERK, promoting PERK autophosphorylation, which further reduces protein translation. Such interaction is facilitated by elevated [Ca^2+^]i. Our findings constitute a clear example of using ER Ca^2+^ as a tool for organelle signal integration. However, the mechanism for active Ca^2+^ release triggered during the acute phase of ER stress remains unknown.

The molecular machinery for Ca^2+^ handling in the ER is conceptually similar to that mentioned above for the plasma membrane; thus, the steady-state of free [Ca^2+^] within the ER depends on the equilibrium between Ca^2+^ uptake by pumps (e.g. SERCA2b), passive Ca^2+^ efflux and buffering by luminal Ca^2+^-binding proteins. The passive leak is a relatively slow process and can be unmasked by application of thapsigargin, a specific inhibitor of SERCAs ^19,20^. At luminal [Ca^2+^] below ∼ 40 µM, basal Ca^2+^ efflux rate is constant and linear, ^19,21^ indicating saturation of leak channels ^22^.

A substantial proportion of passive Ca^2+^ leak (efflux) from the ER occurs via a protein complex termed the “translocon” ^23-27^, which consists of a core heterotrimeric Sec61 complex (Sec61αβγ) and associated proteins ^28-31^. Sec61α, the largest subunit of the heterodimer, spans the entire ER lipid bilayer and forms the pore of the channel through which synthesized proteins are translocated ^32^. In spite of the inner diameter of the channel pore, which varies from 9-15 Å to 40-60 Å, the ribosome and BiP precisely control the ion permeability barrier ^33-36^. Thus, during nascent protein elongation, this is achieved by tight binding of ribosome to the cytosolic side of the ER membrane. The aqueous pore is also closed to the ER lumen by BiP before (ribosome-free state) and during early stages of translocation, until the nascent chain reaches a length of ∼70 amino acids ^32,34^. Only when translocation is completed, the polypeptide chain is released, and ribosome dissociated from ER membrane, is Ca^2+^ ion permeability increased, accounting for reported values of basal Ca^2+^ leak ^23-27^.

We describe here a previously unknown active mechanism for stressor-mediated Ca^2+^ release from the ER. We provide pharmacological and molecular evidence that the translocon generates local [Ca^2+^] increase, particularly during the acute phase of the UPR. When BiP is competitively titrated by misfolded proteins and dissociated from Sec61α translocon, Ca^2+^ efflux is enhanced into cytosol, where it is eventually buffered. Identification of this new Ca^2+^ signal could initiate a paradigm shift in the field by demonstrating that an ER stressor and not a messenger can mobilize Ca^2+^ from the ER.

## Results

### Global and local Ca^2+^ signalling induced by the ER stressor Tunicamycin

Elucidation of Ca^2+^ signalling that triggers and/or regulates the initial phase of the UPR is important in view of the increasingly clear associations of ER stress with numerous pathological processes. Under physiological conditions, Ca^2+^ leak through the translocon is small and does not increase [Ca^2+^]i sufficiently to generate a signal ^20^. In contrast, accumulated ER protein misfolding may give rise to a Ca^2+^ release across translocon channel that mediates early Ca^2+^ signalling in UPR.

We investigated this possible mechanism by directly measuring [Ca^2+^]i in microdomains near ER membrane. For this, we generated a genetically encoded Ca^2+^ indicator, GECI (GCamP6m) attached to the ER membrane ^37^. GCamP6m was fused to 76 carboxyl amino acids of cytochrome b5 corresponding to central region and C-terminal ER-targeting domains (GCamP6-Cytb5). Ten amino acid residues at the C-terminal end are necessary to target the ER membrane, and the next amino acid residues function as a hinge region that increases GECI flexibility ^38^ (Fig. 1a). Importantly, cytochrome b5 is a typical tail-anchored protein located on ER outer surface, and its insertion into the membrane occurs post-translationally and does not involve the Sec61 complex ^39^. The ability of GCamP6-Cytb5 to detect Ca^2+^ microdomains near the ER membrane is greater than that of the cytosolic form of this GECI.

**Figure 1:**
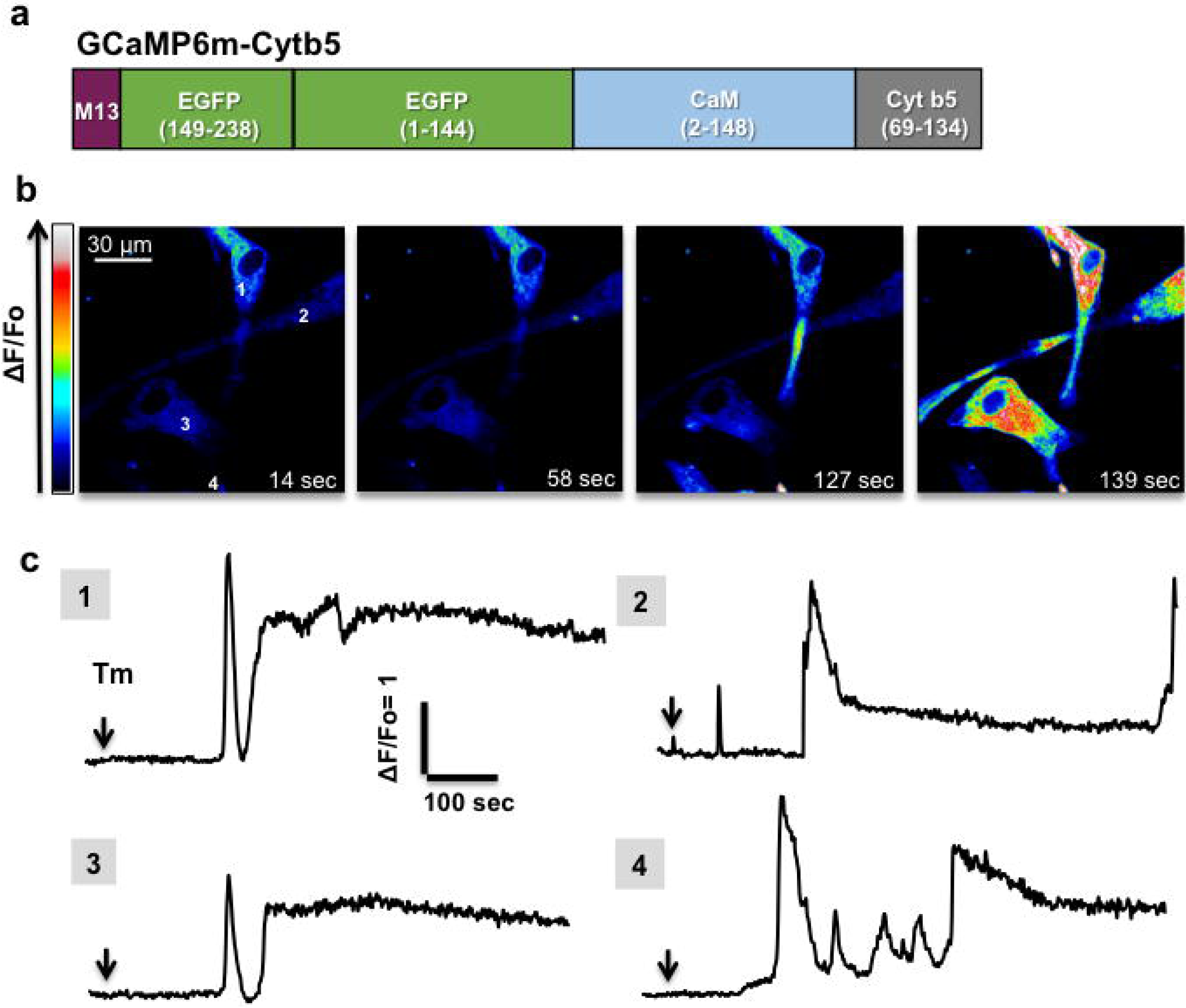
Global calcium increases following addition of high-concentration tunicamycin. Measurement of cytosolic Ca^2+^ changes by confocal imaging of cultured human astrocytes expressing Ca^2+^ indicator GCaMP6-Cytb5. **(a)** Schematic representation of GCaMP6-Cytb5. GCaMP6 was fused at the C-terminus to the transmembrane domain of cytochrome b5 (Cytb5) for tethering to ER membrane. Images were obtained by Nikon Swept Field confocal microscopy (see Methods). **(b)** Single confocal image frames in pseudo-colour showing Ca^2+^ release before and after Tm treatment (Tm; 2.5 µg/ml) at indicated times. Scale bar: 30 µm. Intensity scale bar for these images is shown. **(c)** Fluorescence intensity values were obtained by selecting a 5×5-pixel region from subsequent images during recording of individual astrocytes. These values were normalized against values obtained prior to Tm treatment (ΔF/F_0_), and plotted as a function of time. Numbers 1-4 corresponds to specific regions marked on frame shown in panel (a). A representative experiment from 3 independent experiments is shown.

For induction of ER stress, we used tunicamycin (Tm), whose mechanism of action is different from that of the Ca^2+^ pump inhibitor, thapsigargin. Tm inhibits glycosylation of nascent proteins, resulting in accumulation of misfolded proteins in the ER. Tm was applied in a solution containing only a trace amount of Ca^2+^, such that [Ca^2+^]i increases were due solely to Ca^2+^ release from the ER. Application of a high Tm concentration (2.5 µg/ml) induced Ca^2+^ release that was initiated at one or several sites and propagated the length of the cell. Most of the Ca^2+^ increases were transient, but also induced global Ca^2+^ increases, which did not decay to baseline during the recording period (Fig. 1; Table 1). However, lower Tm concentrations (0.25 - 0.5 µg/ml) inhibited these Ca^2+^ waves (Fig. 2; Table 1). More precisely, reduction of the Tm concentration significantly increased the proportion of astrocytes that displayed Ca^2+^ increases with a narrow spatial spread. On the basis of this spatial spread, we defined localised Ca^2+^ events as either microdomains (spots) (area 1-6 µm^2^) or local areas limited to a part of the cell (area >6-30 µm^2^) (Figs. 1, 2; Table 1). Fluorescence changes were assigned as microdomains when they increased ≥2 SD relative to baseline fluorescence. The mean spatial spreads (full surface at maximal amplitude) were 2.45 ± 0.15 µm^2^ (n=119) for microdomains and 15.82 ±?1.12 µm^2^ (n=33) for local areas. These events constituted the initiation site of a Ca^2+^ wave in ∼26% of cells (n=109), although the actual percentage was likely higher since detection was limited by the position of the focal plane relative to the event. Spontaneous Ca^2+^ microdomains were rarely observed under non-stressed conditions.

**Table 1.**
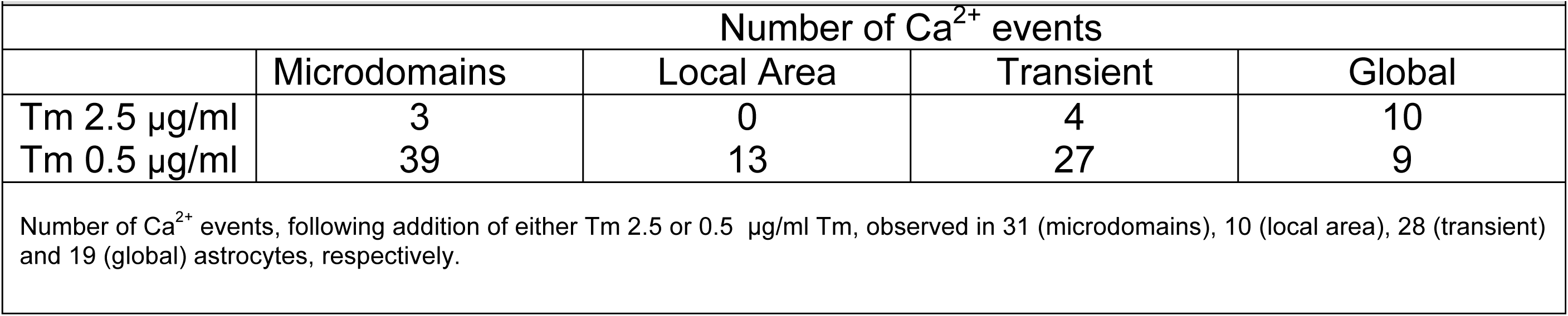
Tunicamycin-induced Ca^2+^ release in astrocytes.

| Number of Ca<sup>2+</sup> events |  |  |  |  |
| --- | --- | --- | --- | --- |
|  | Microdomains | Local Area | Transient | Global |
| Tm 2.5 µg/ml | 3 | 0 | 4 | 10 |
| Tm 0.5 µg/ml | 39 | 13 | 27 | 9 |
| Number of Ca<sup>2+</sup> events, following addition of either Tm 2.5 or 0.5 µg/ml Tm, observed in 31 (microdomains), 10 (local area), 28 (transient) and 19 (global) astrocytes, respectively. |  |  |  |  |

**Figure 2:**
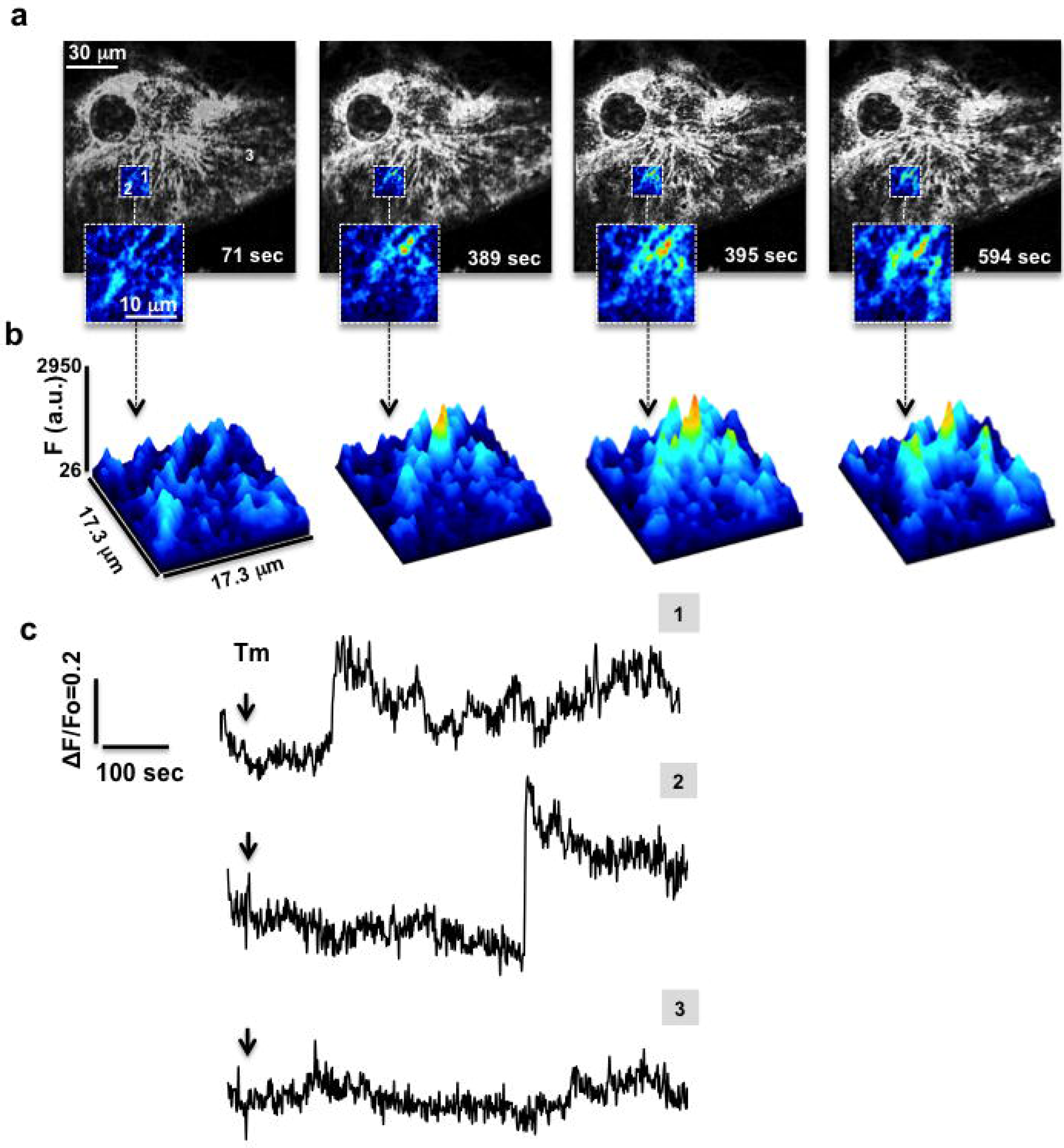
Low-concentration tunicamycin induces several local calcium releases in microdomains near ER. Cytosolic Ca^2+^ changes were measured as in Fig. 1. **(a)** Confocal images (grayscale) corresponding to Ca^2+^ release before (71 sec) and after (389, 395, 594 sec) Tm (0.5 µg/ml) treatment. Insets: magnification of regions with changes in cytosolic Ca^2+^ shown in pseudo-colour. Scale bar: 30 µm. **(b)** Surface plot corresponding to magnified regions, illustrating local changes in cytosolic Ca^2+^. **(c)** Fluorescence intensity values were obtained as in Fig. 1, and ΔF/F_0_ was plotted as a function of time. A representative experiment from 6 independent experiments is shown.

Kinetic characteristics of various patterns of Tm-induced Ca^2+^ release were analysed. We found that the amplitudes of these Ca^2+^ events differed significantly (Table 2). Additionally, the rise times and decay times of Ca^2+^ microdomains were significantly slower than those of local Ca^2+^ areas (Table 2). This phenomenon may be attributable to differing channel compositions of microdomains vs. local areas; *i*.*e*., microdomains have only Tm-mediated slow Ca^2+^ release channels, whereas local areas have fast-release channels as well.

**Table 2.** Analysis of tunicamycin-induced Ca^2+^ release in astrocytes.

| | Astrocytes per group (N) | Amplitude ($\Delta F/F_0$) | Peak Time (sec) | Decay Time (sec) |
| --- | --- | --- | --- | --- |
| Microdomains | 31 | $0.26 \pm 0.018^{* \&}$ | $64.81 \pm 10.643$ | $63.514 \pm 10.042$ |
| Local area | 10 | $0.51 \pm 0.136^{\pi}$ | $24.79 \pm 7.922$ | $36.69 \pm 10.398$ |
| Transient | 28 | $1.20 \pm 0.178^{\diamond}$ | $40.47 \pm 5.663$ | $52.50 \pm 9.491$ |
| Global | 19 | $2.16 \pm 0.423$ | $34.11 \pm 6.743$ | n.d. |

N: number of astrocytes in each category. Mean ± SEM. Statistical significance of differences was analyzed by one-way ANOVA, followed by Tukey's all pair comparison test.
\* Microdomains amplitude differs from transients amplitude ( $P < 0.001$ )
& Microdomains amplitude differs from global amplitude ( $P < 0.0001$ )
$\pi$ Local area amplitude differs from global amplitude ( $P < 0.001$ )
$\diamond$ Transient amplitude differs from global amplitude ( $P < 0.05$ )

A substantial percentage (36.69%; n=109) of Ca^2+^ microdomains (spots) displayed periodic episodes, with a mean frequency 1.21 ± 0.107 events per 50 sec (n=40). Interspike intervals ranged from 21.75 to 100.6 sec, with a mean of 52.43 ± 3.67 sec (n=40). Although a large proportion of Ca^2+^ microdomains displayed periodic episodes, the highest number of occurrences for a single microdomain was six. Of note, application of thapsigargin, which irreversibly inhibits SERCA2b, abolished the periodic episodes (n=11), suggesting that the pump mediates removal of Ca^2+^ microdomains, allowing another round of Ca^2+^ release.

### Tm-induced Ca^2+^ microdomains are regulated by reagents that modify translocon activity

To investigate the possible involvement of translocon channels in localised Tm-induced Ca^2+^ release, we used pharmacological tools that modify translocon permeability. We first examined the effect of emetine, which plugs the translocon by irreversibly inhibiting translational elongation ^40^. Astrocytes expressing ER-anchored GCamP6-Cytb5 were pre-incubated with 100 µM emetine for 30 min and imaged. Emetine blocked the ability of Tm to induce global Ca^2+^ increase (Fig. 3a-c), and resulted in a percentage of cells showing Ca^2+^ microdomains (20.0%, n=20) much lower than control values (45.9%, n=37). Emetine also significantly reduced the amplitude of localised Ca^2+^ increase. In contrast, Tm-induced Ca^2+^ release was enhanced by a 30 min pre-treatment with AB_5_ subtilase (1 µg.ml^−1^), a toxin that specifically cleaves and inactivates BiP ^41^, resulting in a much higher percentage of cells showing global Ca^2+^ increase (29.4%; n=17) relative to controls (5.4%; n=37). This finding was consistent with the increased peak amplitude of the resulting global signal (Fig. 3a-c).

**Figure 3:**
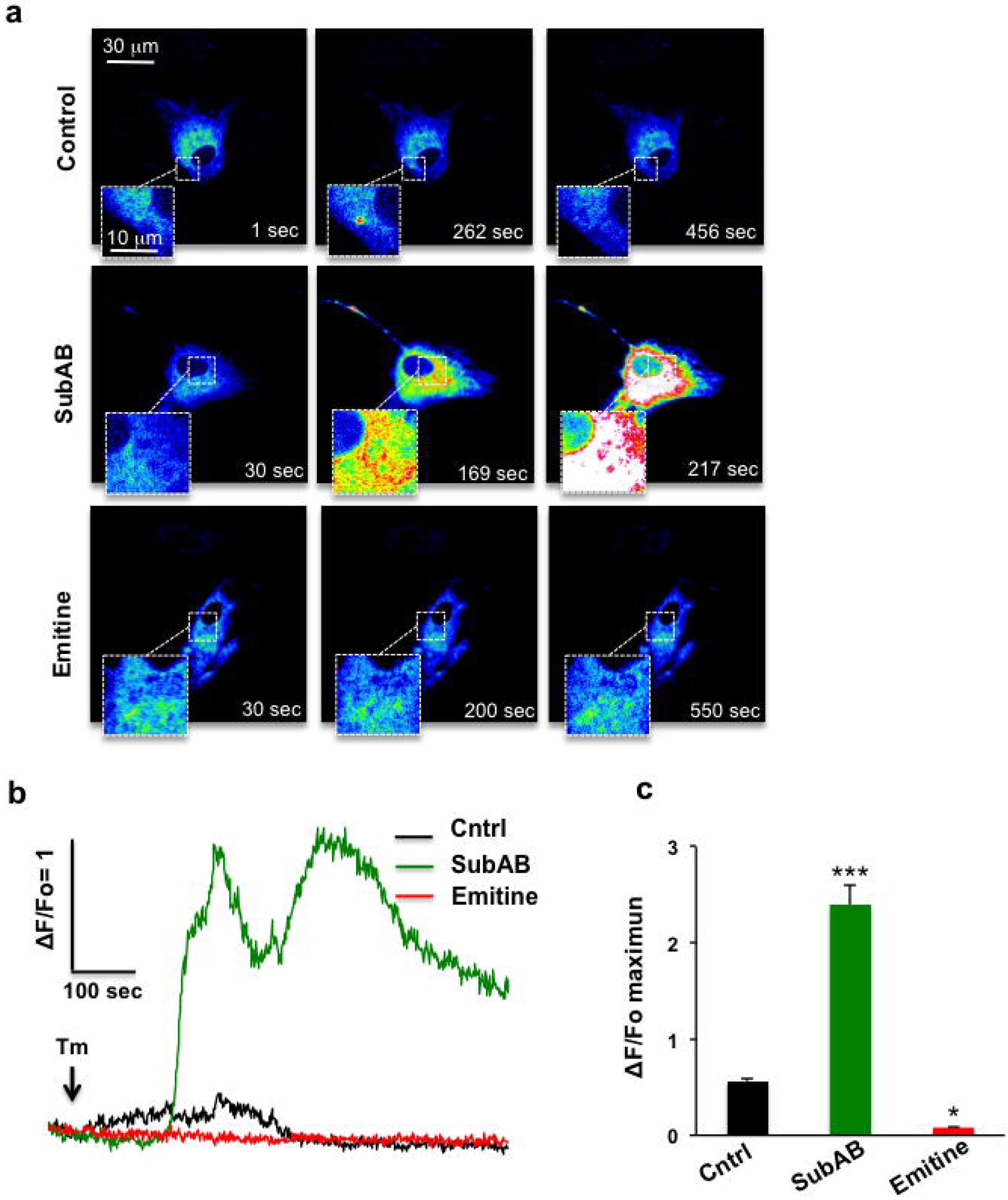
AB_5_ subtilase cytotoxin and emetine modulate Tm-induced local Ca^2+^ increase. Human astrocytes were pre-incubated with either AB_5_ subtilase cytotoxin (SubAB; 1 µg/ml, 30 min) or emetine (1 µM, 30 min), and added with 0.5 µg/ml Tm. **(a)** Sequential confocal images in pseudo-colour illustrating Tm-induced Ca^2+^ release by control, SubAB-treated, and emetine-treated cells. Insets show Ca^2+^ increase events. Scale bar: 30 µm. **(b)** Data for GCamP6-Cytb5 Ca^2+^ increase in terms of ΔF/F_0_ were obtained as in Fig. 1 and plotted as a function of time for each condition. Representative data from 4 independent experiments are shown. **(c)** Histograms (mean ± SEM) showing maximal ΔF/F_0_ for each condition. *p< 0.05, ***p< 0.0001 (ANOVA, Tukey’s HSD test).

We next examined the effect of anisomycin, which inhibits elongation by locking nascent chains in the ribosome ^42^, on Tm-induced Ca^2+^ release. Anisomycin pre-treatment (60 min; 200 µM) completely abolished Tm-induced global Ca^2+^ increase, and greatly reduced the percentage of cells showing local Ca^2+^ increase (16.6%, n=6; relative to 45.9%, n=37 for control) (Fig. S1). Tm-induced Ca^2+^ release was amplified by puromycin, an antibiotic that purges translocons from nascent polypeptide chains ^43^ (Fig. S1). The percentage of cells showing global Ca^2+^ increase was much higher for those treated with 20 µM puromycin (40.0%, n=6) relative to controls (5.4%, n=37). It should be note that this effect was observed only when puromycin and anisomycin were added at the same time. Pre-incubation with puromycin inhibited Tm-induced Ca^2+^ release; the antibiotic was likely able to induce Ca^2+^ leak and deplete the ER of Ca^2+^ within a few minutes. These findings, taken together, provide pharmacological evidence that the translocon channel *per se* mediates Tm induction of Ca^2+^ microdomains.

### Subdivision of Tm-induced Ca^2+^ responses into local events

The dynamics of Tm-induced Ca^2+^ microdomains were further analysed using either a slow Ca^2+^ buffer EGTA or IP_3_ and ryanodine receptor inhibitors (xestospongin 3 µM/ ryanodine 50 µM; “xesto/ryano”). Both of these treatments significantly increased the percentage of astrocytes displaying Tm-induced Ca^2+^ microdomains (Fig. 4a). Activity was induced in 16.40 ± 0.2% (n=6) of EGTA-treated and 83.50 ± 16.5% (n=5) of xesto/ryano-treated cells, but in only 12.16 ± 0.6% (n=8) of control cells.

**Figure 4:**
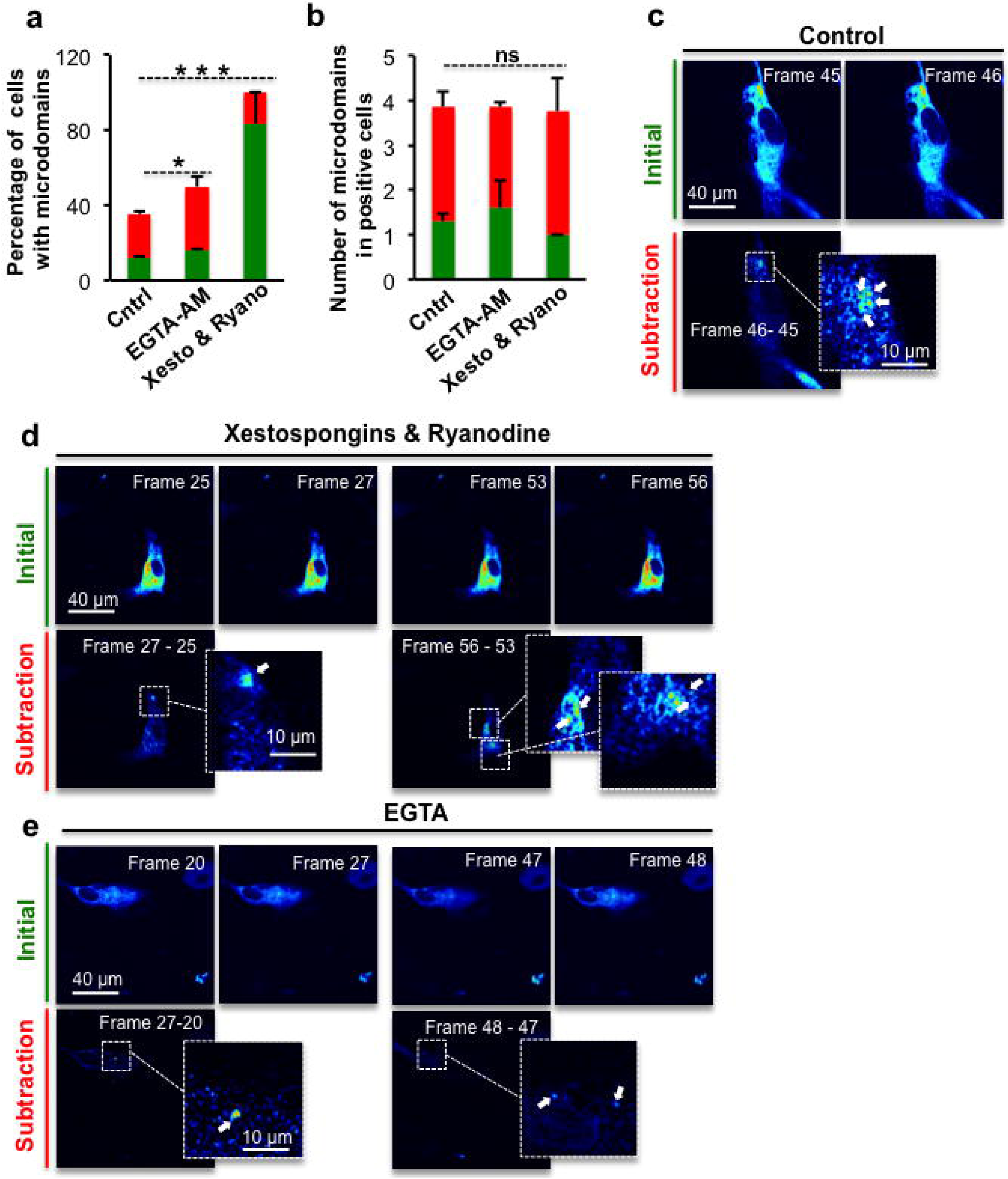
EGTA-AM and xesto/ryano treatments increase the likelihood of local Ca^2+^ increase following Tm addition. Human astrocytes were pre-incubated with either EGTA-AM (1 µM, 20 min) or xesto (3 µM, 30 min)/ ryano (50 µM, 60 min), and added with 0.5 µg/ml Tm. **(a)** Sequential subtraction: pixel-by-pixel intensity values for each frame were subtracted from values of some frames ahead for clear visualisation of microdomains. Percentages of cells with microdomains before and after subtraction (mean ± SEM) for each condition are shown respectively as red and green. **(b)** Total numbers of Ca^2+^ microdomains before and after subtraction in positive cells (mean ± SEM) for each condition are shown respectively as red and green. **(c-e)** Confocal image stacks in pseudo-colour before and after subtraction for control **(c)**, xesto/ryano-treated **(d)**, and EGTA-treated **(e)** cells. Insets show microdomains. Scale bar: 40 µm (regular images) or 10 µm (magnified images). Representative data from 5 independent experiments are shown. *p< 0.05, ***p< 0.0001, ns: not significant (ANOVA, Tukey’s HSD test).

A sequential subtraction process was performed on the entire image sequence to help detect the localised Ca^2+^ increases induced by Tm. Thus, pixel-by-pixel fluorescence intensity values for each frame were subtracted from values of the image a few frames ahead (Fig. 4c-e). This analysis revealed a further increase in the percentage of responding cells by providing clear visualization of new Ca^2+^ spots, particularly after EGTA and xesto/ryano treatments. Greater temporal resolution would presumably also reveal an increased incidence of Tm-induced Ca^2+^ microdomains in control cells (Fig. 4a). However, for positive cells, microdomain numbers did not differ between the conditions tested (Fig. 4b).

In contrast to the clear increase in proportions of cells showing Ca^2+^ microdomains under these treatments, the percentages of cells showing broader events, such as local areas of Ca^2+^, were smaller after xesto/ryano (16.6%, n=5) or EGTA treatment (11.1%, n=10) relative to controls (60%, n=17).

These findings suggest that EGTA and xesto/ryano subdivide the Ca^2+^ response into localised events that inhibit wave propagation, and refine the spatio-temporal profile of Ca^2+^ microdomains. Ca^2+^ spots are considered to represent a spatial-temporal summation of a single ion channel activated by CICR. The spatial spread at the time of peak amplitude of Tm-induced microdomains following xesto/ryano treatment (2.51 ± 0.31 µm^2^, n=13) was not significantly different from that observed in control cells (2.58 ± 0.20 µm^2^, n=27), indicating that the microdomains are constituted only by translocons. In contrast, spatial spread was significantly narrower (1.65 ± 0.24 µm^2^, n=19; p< 0.05 by ANOVA) for EGTA-treated cells, indicating an active CICR mechanism within each translocon cluster.

### Tm-induced Ca^2+^ signalling is independent of IP_3_ Receptors

Properties of Ca^2+^ events generated by the translocon were further elucidated by experiments using Human Embryonic Kidney cells (HEK-293) in which all three IP_3_R isoforms were knocked out (TKO-HEK); these cells do not express ryanodine receptors. High Tm concentration (2.5 µg/ml) induced discrete, highly localised, transient Ca^2+^ microdomains, lasting an average of 465 ± 21.14 sec (n=10), in all TKO-HEK cells (Fig. 5). This activity was completely abolished by pre-incubation with 100 µM emetine (n=3).

**Figure 5:**
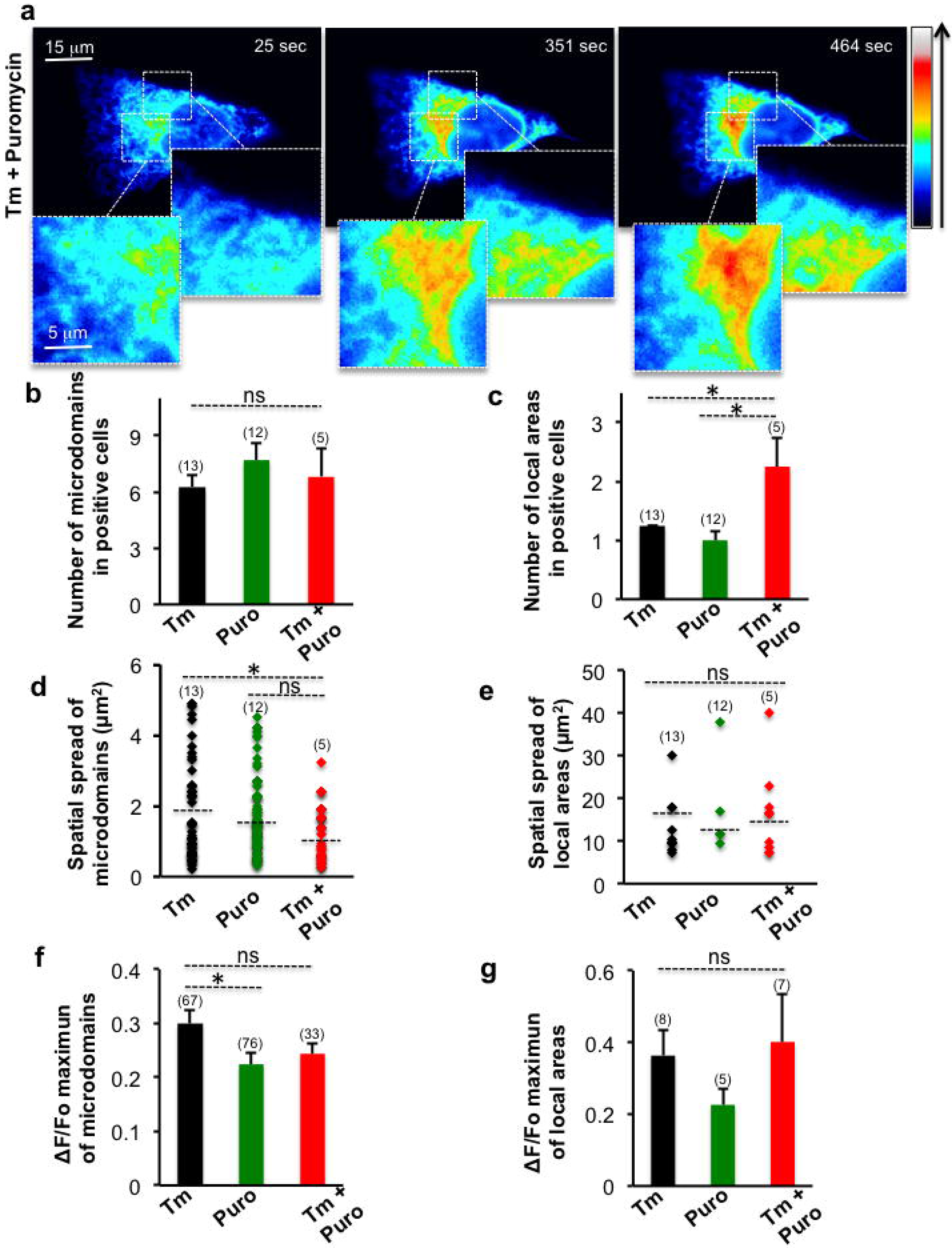
Puromycin treatment enhances Tm-induced local Ca^2+^ increase in TKO-HEK cells. Human Embryonic Kidney cells (HEK-293) with knockout of all three IP_3_R isoforms (termed TKO-HEK) were treated with Tm (2.5 µg/ml) and puromycin (20 µM). Images were obtained by Olympus Spinning Disk confocal microscopy (see Methods). **(a)** Confocal images in pseudo-colour illustrating Ca^2+^ release in response to Tm + puromycin addition. Scale bar: 15 µm (regular image) or 5 µm (magnified image). **(b-c)** Histograms (mean ± SEM) showing numbers of microdomains **(b)** or local areas **(c)** in positive cells for each condition. **(d-e)** Dot plots (mean ± SEM) showing spatial spread (µm^2^) of microdomains **(d)** or local areas **(e)** for each condition. **(f-g)** Histograms (mean ± SEM) showing maximal ΔF/F_0_ of microdomains **(f)** or local areas **(g)** for each condition. *p< 0.05, ns: not significant (ANOVA, Tukey’s HSD test).

The Tm-induced Ca^2+^ signal was propagated as a local area of Ca^2+^ in 73% of TKO-HEK cells (n=10), without triggering global Ca^2+^ waves. The percentage of cells displaying localized Ca^2+^ areas was even higher (80%, n=5) following a combination treatment with Tm and puromycin (Fig. 5c). Tm-induced Ca^2+^ release was enhanced by this treatment combination, resulting in a number of local Ca^2+^ areas much higher than that in cells treated with Tm or puromycin alone (Fig. 5a,c; Fig. S2). The numbers of microdomains per cell did not differ significantly among these treatments (Fig. 5b).

Another interesting feature of Ca^2+^ signaling in TKO-HEK cells is that the spatial spread at the peak amplitude time of Tm+puromycin-induced microdomains was much narrower than that of Tm-induced microdomains (Fig. 5d). This finding is consistent with the enhanced Ca^2+^ activity as evidenced by the higher number of local Ca^2+^ areas observed in Tm+puromycin-treated cells (Fig. 5c). This treatment clearly facilitates propagation of Ca^2+^ microdomains and consequent formation of new local Ca^2+^ areas. Local Ca^2+^ areas under the three treatments did not differ significantly in mean spatial spread or fluorescence amplitude (Fig. 5e-g), suggesting a self-limited regulation of these Ca^2+^ events.

We conclude from these data that cross talk of Ca^2+^ release between clusters strongly enhances cell excitability under conditions of ER stress, even though the translocon is able to induce only localised Ca^2+^ events.

### BiP expression regulates Tm-induced Ca^2+^ release

It was previously demonstrated that BiP played a role in sealing Sec61 pores ^34,36^. To examine the possible role of BiP in triggering Ca^2+^ release through the translocon under ER stress, we overexpressed BiP in TKO-HEK cells and performed confocal Ca^2+^ imaging. Red fluorescence protein mCherry was fused to BiP C-terminal domain, followed by the ER-retrieval motif (amino acid residues KDEL). Of note, cells with co-overexpression of GCamP6-Cytb5 and BiP-mCherry showed a typical ER network with green and red fluorescence proteins, respectively (Fig. 6a).

**Figure 6:**
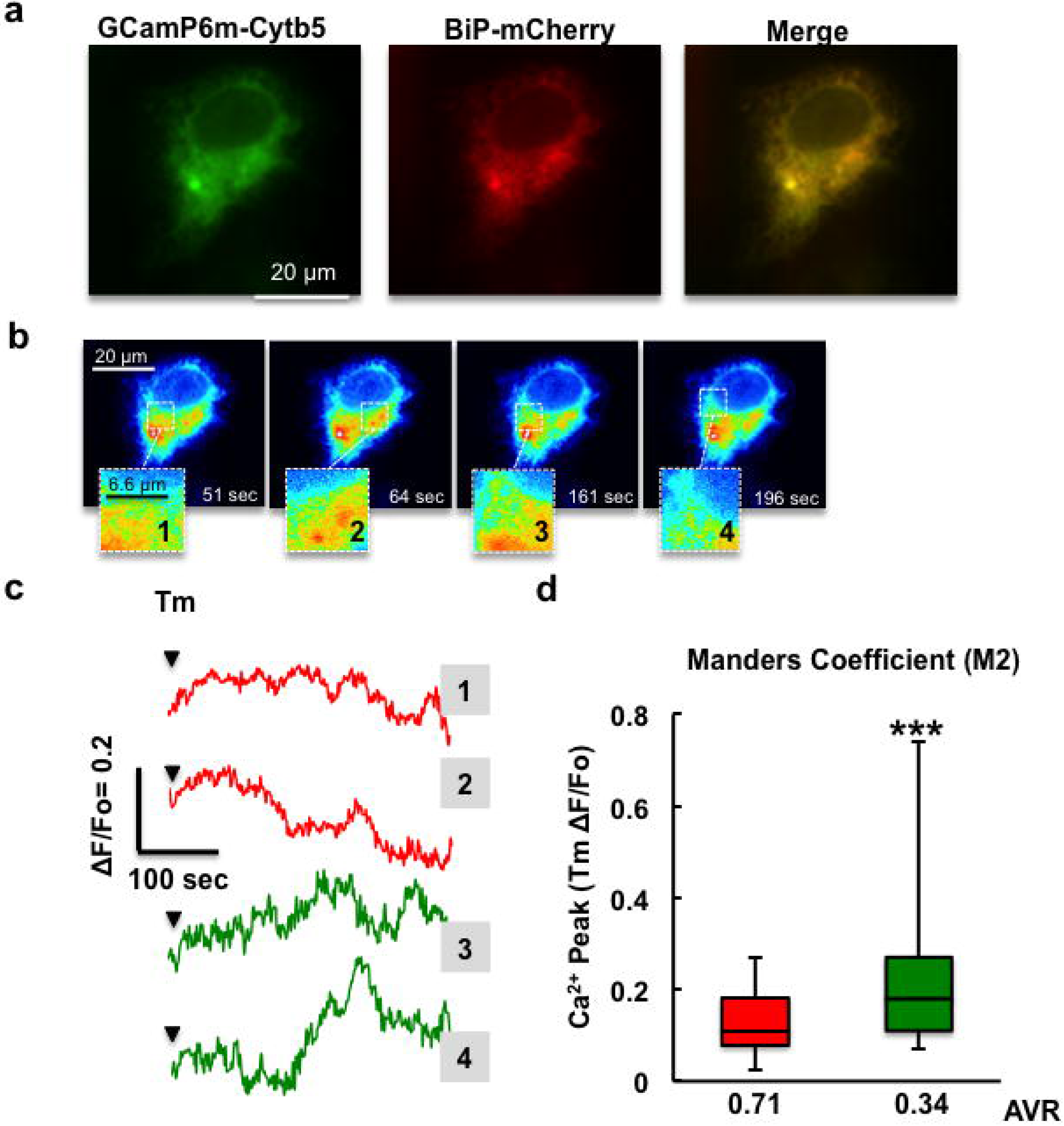
Tm-induced local Ca^2+^ increase is reduced by chaperone mCherry-BiP overexpression. TKO-HEK cells co-overexpressing either GCaMP6-Cytb5 and mCherry-BiP or GCaMP6-Cytb5 and empty mCherry vector were added with 2.5 µg/ml Tm, and Ca^2+^ confocal images were taken. **(a)** Representative images of GCamp6-Cytb5 (green) and mCherry-BiP (red) expression by TKO-HEK. Scale bar: 20 µm. **(b)** Confocal image sequences in pseudo-colour illustrating Tm-induced Ca^2+^ release for each condition. Insets show Ca^2+^ increase events. Scale bar: 20 µm. The two fluorophores were recorded alternately. **(c)** Data for GCamP6-Cytb5 Ca^2+^ increase in terms of ΔF/F_0_ were obtained as in Fig. 1 and plotted as a function of time for each condition. Representative data from 3 independent experiments are shown. **(d)** MOCs were calculated for a 5×5-pixel ROI showing Ca^2+^ release. MOC **M2** was pooled as sets of values <0.5 and >0.5 (low and high co-localisation, respectively), and average. M2 averages were plotted vs. average peak Ca^2+^ responses. Data were pooled from 3 independent experiments. ***p< 0.0001 (ANOVA, Tukey’s HSD test). ROIs in which mCherry-BiP overlapped GCamP6-Cytb5 displayed a significant reduction in amplitude of Ca^2+^ release (shown as red in (c) and (d)) relative to ROIs with no overlap (shown as green in (c) and (d)).

Application of 2.5 µg/ml Tm induced only Ca^2+^ microdomains (Fig. 6b). We measured GCamP6-Cytb5 fluorescence intensity (ΔF/F_0_) in certain regions of interest (ROIs) and then calculated Mander’s Overlap Coefficient (MOC) for those ROIs. The obtained MOC M2 values with maximum ΔF/F_0_ values were pooled in two sets (MOC M2 <0.5 and >0.5), reflecting low and high co-localisation of GcamP6-Cytb5 and BiP-mCherry. The peak Ca^2+^ amplitude was significantly lower in ROIs in which BiP-mCherry overlapped with GCamP6-Cytb5 (MOC M2 >0.5) compared to those in which BiP expression was low (Fig. 6c,d). As negative control, cells were co-transfected with GCamP6-Cytb5 and empty vector carrying only mCherry. ROIs with high vs. low overlap of mCherry and GCamP6-Cytb5 fluorescence did not differ significantly in Ca^2+^ release (Fig. S3).

These findings show that elevated BiP expression blocks Tm-induced Ca^2+^ release across translocon, suggesting the need for BiP to dissociate from the Sec61 channel in order to allow an increased Ca^2+^ efflux through the translocon. This would naturally occur once ER stress has been activated and BiP titrated by accumulated unfolded protein.

### Local Ca^2+^ events generated at the translocon are inhibited by high Ca^2+^ concentrations

Given that BiP plays a role in gating the luminal end of translocon pore in stressed cells, the next question we asked was what happens to the cytosolic end of translocon pore, and its association with the ribosome, during Tm-induced Ca^2+^ release process? To address this question, we used a combination of two approaches: confocal Ca^2+^ imaging in living cells, and subsequent fluorescence immunocytochemistry. Ca^2+^ imaging was performed in TKO-HEK cells expressing GCamP6-TMCytb5 following Tm and puromycin addition; immediately after that Ca^2+^ release was detected, cells were rapidly fixed in formaldehyde (Fig. 7a-c). Following confirmation that GCamP6 fluorescence was not completely quenched by fixation, cells were labelled with anti-Sec61α and anti-S6-ribosomal protein antibodies and visualized using fluorescence-conjugated secondary antibodies (Fig. 7e). Dishes with imprinted grids were used for identification of each fixed and immunostained cell recorded after Ca^2+^ imaging (Fig. 7c), and then confocal imaged (Fig. 7d).

**Figure 7:**
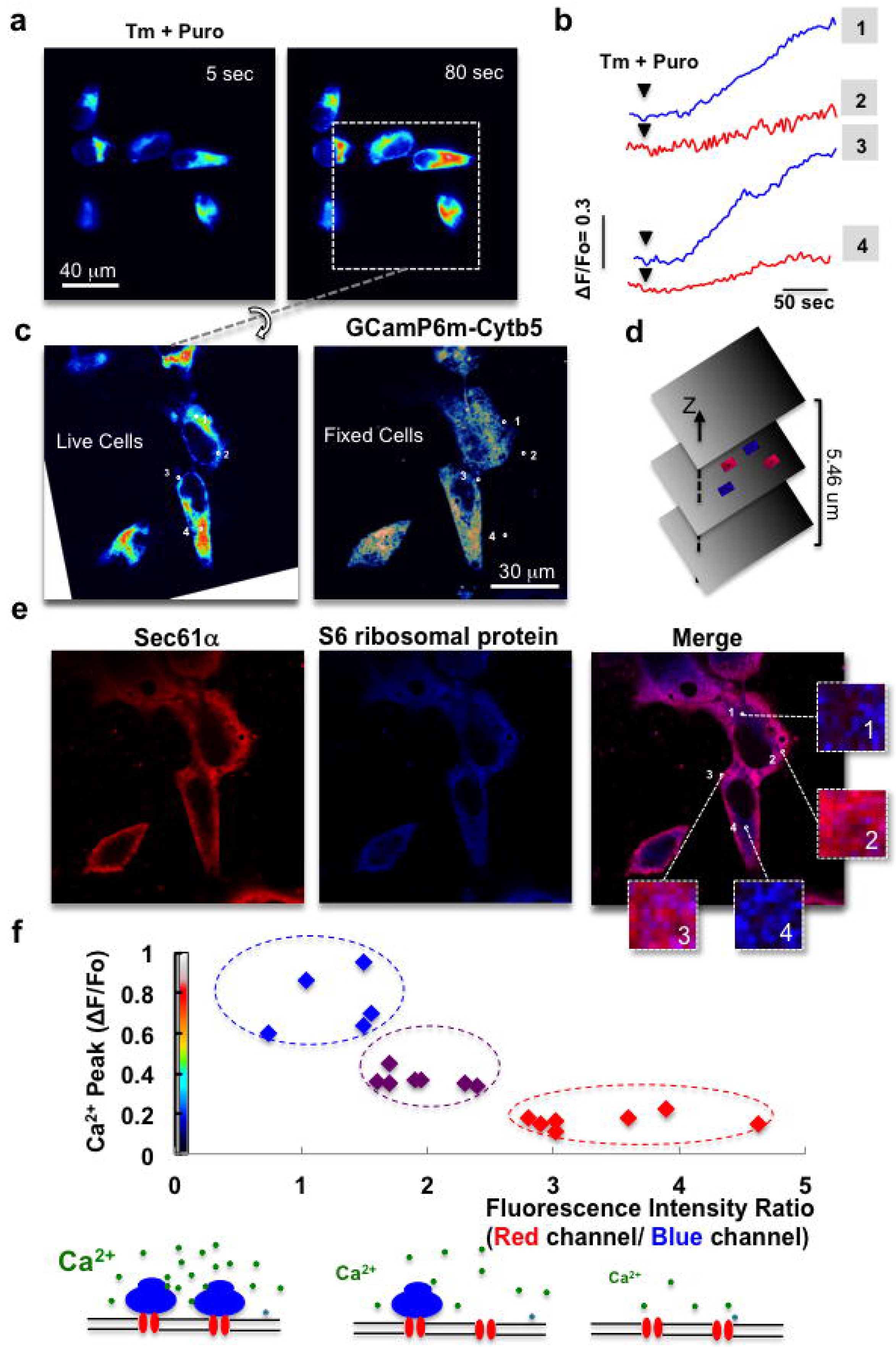
Cytosolic Ca^2+^ concentration regulates Tm-induced local Ca^2+^ increase. TKO-HEK cells expressing GCamP6-Cytb5 cultured in dishes with imprinted grids were added with Tm (0.5 µg/ml) and puromycin (20 µM). **(a)** Confocal image sequences in pseudo-colour illustrating Tm-induced Ca^2+^ release before (5 sec) and after (80 sec) Tm + puromycin treatment. Scale bar: 40 µm. **(b)** Data for GCamP6-Cytb5 Ca^2+^ increase in terms of ΔF/F_0_ were obtained as in Fig. 1 and plotted as a function of time for each cell. Representative data 3 independent experiments are shown. **(c)** Cells were fixed in formaldehyde immediately after Ca^2+^ release was detected. Left: last stack of Ca^2+^ imaging recorded. Right: same group of cells after fixation. Each fixed cell recorded in Ca^2+^ imaging was identified and then rotated. Numbers 1-4 indicate particular regions with differing magnitudes of Ca^2+^ increase (see (b)). **(d, e)** Fixed cells were immunostained with anti-S6 ribosomal protein or anti-Sec61α primary antibodies and with Alexa-conjugated anti-rabbit Alexa Fluor 568 and anti-mouse Alexa Fluor 405 (respectively) secondary antibodies. Optical sectioning of confocal images was performed using Zeiss LSM 800 confocal microscope. **(d)** Schematic representation of 21 optical sections (plane thickness: 0.23 µm; total thickness: 5.46 µm). Fluorescence intensities were analysed only in the Z-plane, in which blue and red regions were obvious. **(e)** Fixed cells were stained with anti-S6 ribosomal protein (shown as red), anti-Sec61α (shown as blue), antibodies, or corresponding merge. Magnified image shows ROI in which Ca^2 +^ was measured (see (b)). There are clear immunostaining differences between ROIs #1 and #4 vs. #2 and #3. **(f)** Red and blue fluorescence intensities were expressed as a ratio and plotted vs. changes in GCamP6-Cytb5 fluorescence (ΔF/Fo) for the same ROIs. Data were pooled from 3 independent experiments. ROIs in which ribosome-free translocons were abundant displayed a significant reduction in amplitude of Ca^2+^ release (shown as red in (f)) relative to ROIs with predominance of ribosome-bound translocons (shown as blue in (f)).

Immunostaining of the Sec61α pore has a well-known peculiarity: antibody access to translocon channel epitopes in cross-linked cells is sterically obstructed by the large ribosome structure ^34,44^. As consequence, cellular regions (ROIs) in which ribosome-bound translocon predominates are clearly distinct from ROIs in which ribosome-free translocons are abundant. We therefore analysed fluorescence intensities only in the Z-plane in which distinct ROIs with red (Sec61α) and blue (S6-ribosomal protein) fluorescence were obvious, and presumably the ER was most abundant (Fig. 7d). The degree of anti-Sec61α antibody labelling in stressed and fixed TKO-HEK cells was not uniform (Fig. 7e). The degree of anti-S6-ribosomal protein antibody labelling was highest in ROIs with little or no Sec61α fluorescence, consistent with the steric blockage caused by ribosomal structure.

The ratio of red to blue fluorescence intensity was plotted vs. the change of GCamP6 fluorescence (ΔF/Fo) in the same ROIs (Fig. 7f), and a functional correlation of this ratio with amplitude of Ca^2+^ release was observed. Surprisingly, ROIs with predominant red fluorescence, indicating greater abundance of ribosome-free translocons, had low Ca^2+^ release. Lower red/blue ratio was found for ROIs with intermediate Ca^2+^ release. ROIs with high Ca^2+^ release had predominantly blue fluorescence, indicating an abundance of ribosome-bound translocons. We note that the latter group of ROIs features two underlying processes: maximal Ca^2+^ release across translocon, which is likely followed by the subsequent binding of the ribosome to Sec61 pore that inhibits further Ca^2+^ release. Previous reports suggest inhibition may be mediated by a Ca^2+^-binding protein ^45-47^.

Taken together, the findings suggest [Ca^2+^]i differentially regulates Ca^2+^ release through translocon under ER stress. [Ca^2+^]i can be either stimulatory or inhibitory, depending on the concentration. This dual modulation of Ca^2+^ release is typical of ion channels ^48-51^.

## Discussion

This study focused on the translocon as new Ca^2+^ signalling system in the context of the early unfolded protein response (UPR). To our knowledge, local ER Ca^2+^ dynamics linked directly to ER stress have not been previously investigated.

The translocon has previously been reported to function as a passive Ca^2+^ leak channel that counteracts Ca^2+^ uptake via SERCA2b pump ^23-27^. Other putative basal Ca^2+^ leak channels in ER, involved in some way in UPR, include Bax inhibitor-1 ^52,53^, BCL-2/IP_3_R ^54,55^, presenilins ^56^, and ryanodine receptors in cardiac cells ^57^. In addition, TMCO1 has been described as an active Ca^2+^ channel that undergoes oligomerisation in response to Ca^2+^ overloading in ER lumen ^58^.

None of the above studies directly monitored [Ca^2+^]i changes attributable to a specific leak channel. We utilized an ER membrane-tethered form of a genetically encoded Ca^2+^ indicator (GCamP6-Cytb5), which overcomes the limitations of known cytosolic forms of chemical or genetically encoded Ca^2+^ indicators. This approach allowed detection of highly localised Ca^2+^ microdomains within ER-stressed cells, thereby permitting us to evaluate the new Ca^2+^ signal.

Generation of Ca^2+^ microdomains immediately following initiation of the UPR appears to be based on two processes. First, BiP is dissociated from luminal domain of Sec61α when it is titrated by unfolded protein to promote luminal protein folding. As a consequence, Tm-induced Ca^2+^ events were amplified by pre-incubation with SubAB cytotoxin (which specifically hydrolyses BiP), and were inhibited by BiP overexpression. Secondarily, the reduction of protein synthesis is also followed by detachment of ribosome from translocon and release of polypeptide chain. Thus, Tm-induced Ca^2+^ microdomains were abolished by either emetine or anisomycin, which locked polypeptide chains in ribosome and Ca^2+^ events were enhanced by puromycin, which induced premature release of nascent polypeptide chains. These two processes jointly permitted release of sufficient Ca^2+^ across translocon to generate a detectable Ca^2+^ signal that was likely further amplified by CICR (Ca^2+^ positive feedback) to increase the range over which neighbouring translocons can be recruited. Loading of EGTA decouples the discrete Ca^2+^ event, thereby inhibiting subsequent global signalling, and also reduced the area of each Ca^2+^ microdomain.

A number of investigators have documented in depth, the underlying mechanisms of on IP_3_-induced Ca^2+^ puffs ^59-61^. In comparison, the properties of Tm-induced Ca^2+^ microdomains appear to be well conserved overall, although several differences were evident. For example, relative to convectional Ca^2+^ puffs in mammalian cells, Tm-induced Ca^2+^ microdomains have significantly slower kinetic parameters (time to peak, and decay times) and lower fluorescence amplitude (ΔF/Fo 0.2, vs. 0.45) ^62^.

The smaller values of these parameters and narrower spatial spread observed for Tm-induced Ca^2+^ microdomains (mean 2.45 µm^2^, vs. ∼7 µm^2^ for IP_3_-induced Ca^2+^ puff) are consistent with the lower Ca^2+^ channel conductance for translocon relative to IP_3_Rc ^62^. In spite of the differences in properties, the total number of Ca^2+^ microdomains per cell in astrocytes is similar to the number of IP_3_ puffs generally reported in mammalian cells (∼ 4) ^62-64^. This may reflect, as suggested by I. Parker’s group for IP_3_-induced puffs ^62^, the localisation of Ca^2+^ events at distances similar to the diffusional range of action of Ca^2+^ in cytosol ^65^, whereby cells can maintain control of the explosively regenerative CICR mechanism to induce appropriate local and global signals. Even the fact that Ca^2+^ microdomains generated by translocons function as initiation sites of subsequent Ca^2+^ waves, under certain conditions, may depend on close proximity of IP_3_Rc. Indeed, for TKO-HEK cells, in which there is no functional coupling between the translocon and IP_3_Rc, Ca^2+^ increase propagating throughout the cell was not observed under any ER stress condition tested. This model system displayed only localised Ca^2+^ events, and in comparison with wild-type mammalian cells, showed notably increased numbers of both Ca^2+^ microdomains and Ca^2+^ local areas. These data suggest an active cross talk between translocon clusters. Our data further suggest an intrinsic Ca^2+^ excitability of translocon, which depends, in turn, on ER Ca^2+^ content, since even brief pre-incubation with puromycin (which reduces Ca^2+^ content) abolishes subsequent Tm-induced Ca^2+^ release. These findings, in conjunction with the observation that BiP overexpression significantly reduced amplitude of Tm-induced Ca^2+^ release, indicate that the ER Stress induced Ca^2+^ signal occurs mainly during the early phase of UPR, prior to BiP upregulation and to depletion of luminal Ca^2+^ content.

We observed an additional level of regulation by [Ca^2+^]i near the translocon channel. [Ca^2+^]i appears to regulate Ca^2+^ efflux across translocon in a biphasic manner; *i*.*e*., its effect is stimulatory at low concentrations and inhibitory at high concentrations. Tm-induced Ca^2+^ microdomains therefore display kinetics typical of ion channels modulated by Ca^2+^, with the activation processes faster than inactivation processes ^48-51^. Regions with low Tm-induced Ca^2+^ release clearly contained ribosome-free Sec61 complex, consistent with open status of translocon channel. In contrast, areas with high Tm-induced Ca^2+^ release had high ribosome density of ribosomes indicating that the Sec61 complex was ribosome-bound, presumably reflecting ion-impermeable channel configuration ^33^. Such dual Ca^2+^ modulation may be mediated by both direct action of Ca^2+^ on the channel and indirect action through a Ca^2+^-binding protein such as calmodulin (CaM) ^45-47^, which binds Ca^2+^ with stoichiometry four and low affinity (Kd ∼10-12 µM) ^66^. Specifically, R. Zimmermann’s group characterized the translocon as a basal Ca^2+^ leak channel and showed that Ca^2+^-CaM bound to conserved IQ motif present in cytosolic domain of Sec61α, thereby limiting Ca^2+^ permeability of the channel by recruiting ribosomes to translocon complex ^67^. Our findings are consistent with the model that CaM also participates in restricting Tm-induced Ca^2+^ release across translocon through ribosome recruitment to the complex. We also note that removal of Ca^2+^ from Ca^2+^ microdomains is presumably mediated by SERCA2b ^68,69^, leading to dissociation of Ca^2+^ from CaM and consequent reversal of its inhibitory effect on Ca^2+^ efflux through the Sec61 channel. This process would initiate another round of Ca^2+^ release and help account for periodic episodes of Tm-induced localised Ca^2+^ events. In fact, blocking of SERCA activity by thapsigargin abolished repetition of local Tm-induced Ca^2+^ events.

In summary, our findings reveal the existence of a novel Ca^2+^ signalling system, initiated by Ca^2+^ microdomains, which is activated during the early phase of UPR. A proposed molecular model is described schematically in Fig. 8, which also takes into account observations regarding CaM modulation of translocon Ca^2+^ leak by Zimmermann’s group ^67^. The close proximity of Tm-induced Ca^2+^ microdomains to ER membrane indicates their essential role in local modulation of UPR components. Future studies will clarify the functional significance of this novel Ca^2+^ signalling system in ER stress processes and cellular responses.

**Figure 8:**
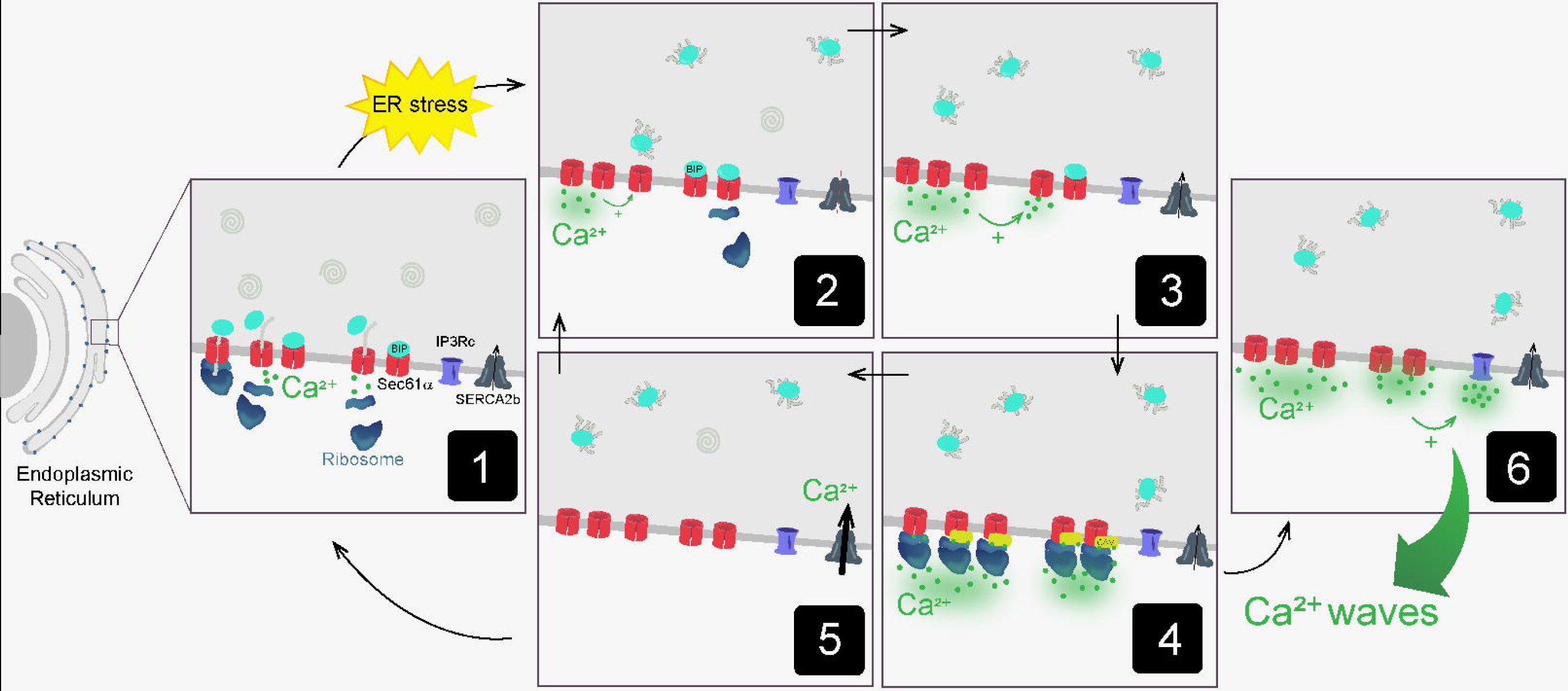
Ca^2+^ signal generated by translocon during early phase of ER stress. **(1)** Steady state of ER: protein (shown as spirals) processing and luminal Ca^2+^ concentration are optimal. Most translocon pores are blocked by BiP and/or ribosomes, maintaining the permeability barrier. When translocation is completed, ion permeability increase as a result of release of nascent chain and dissociation of ribosomes from Sec61 complex, accounting for passive Ca^2+^ leak. SERCA2b counteracts the loss of Ca^2+^. **(2)** ER stress: protein translation is attenuated, and BiP titrated by unfolded protein is dissociated from luminal domain of ribosome-free Sec61α. Ca^2+^ release through translocon is enhanced, and further amplified by CICR, which recruit neighbouring translocons. **(3)** New translocon clusters are activated by Ca^2+^ positive feedback. **(4)** High local Ca^2+^ concentration becomes inhibitory; it binds CaM that engage ribosomes to block translocon Ca^2+^ flux, such that Ca^2+^ signal remains a local event. **(5)** SERCA2b is activated and mediates removal of Ca^2+^ from Ca^2+^ microdomains, and consequent dissociation of Ca^2+^ from CaM. Cessation of Ca^2+^ inhibition accounts in part for generation of repetitive Ca^2+^ microdomains. **(6)** Ca^2+^ released from translocon clusters activates Ca^2+^ flux through IP_3_Rc by explosive CICR mechanism, resulting in generation of Ca^2+^ waves (global signal).

## Methods

### Reagents and antibodies

Reagents and antibodies were from Sigma-Aldrich or Fisher Scientific unless specified otherwise.

### Constructs and plasmids

We used a genetically encoded Ca^2+^ indicator tethered to ER membrane (Custom DNA Constructs; University Heights, OH, USA). GCamP6m cDNA was subcloned into pcDNA3.1 vector, engineered to carry cDNA corresponding to C-terminal 76 amino acid residues of rat cytochrome b5 ^38^, and termed **p**GCamP6m-Cytb5.

mCherry-BiP-KDEL construct was from Addgene (Watertown, MA, USA) (plasmid #62233; **p**Bip-mCherry).

### Human tissues, and cell culture

Human astrocytes were obtained from brain tissues of male patient (age 40 years) and female patients (ages 33, 44, and 50 years) as described by D.T. Lin et al. ^70^. Patients provided informed consent to the Dept. of Neurosurgery, Univ. of Texas Health Science Center, San Antonio, TX, USA, and protocols were approved by the institutional Ethics Committee.

Brain tissues were minced using a sterile razor and trypsinized (trypsin-EDTA 0.25%; #25200056; Life Technologies Corp.; Carlsbad, CA, USA) for 30 min in a 37 °C humidified incubator. Cells were suspended in fresh DMEM/F-12 (#11039-021) supplemented with 10% FBS (#12483-020), 10,000 U/ml penicillin, and 10 mg/ml streptomycin (#15140-122).

Triple IP_3_R knockout human embryonic kidney cell line (TKO-HEK) from Kerafast (Boston, MA, USA; #EUR030) was grown in DMEM (#11995-065) supplemented with 10% FBS, 10,000 U/ml penicillin, and 10 mg/ml streptomycin. All cell cultures were incubated at 37 °C in humid 5% CO_2_ atmosphere.

Cell cultures were tested for the presence of Mycoplasma using PCR-based method.

### Transfections

Human astrocytes or TKO-HEK cells were plated in either 35 mm dishes or P35G-1.5-14-C (MatTek Corp.; Ashland, MA, USA) as indicated, and transfected with 2 µg cDNA **p**GCamP6m-Cytb5 and 2 µl transfection agent X-tremeGENE (#06366244001) as per manufacturer’s instructions. Calcium imaging was performed either three or one days after transfection, respectively, for the two types of cells.

TKO-HEK cells were co-transfected with 2 µg plasmid corresponding to either **p**Bip-mCherry, or empty vector and **p**GCamP6m-Cytb5, as per manufacturer’s instructions. [Ca^2+^]i changes were recorded one day later.

### Ca^2+^ imaging

Culture medium was replaced by low-Ca^2+^ buffer (in mM: 15 HEPES/ NaOH, 130 NaCl, 5.4 KCl, 2 MgCl_2_, 10 glucose). Cytosolic calcium imaging was performed: (i) for human astrocytes, using a Nikon Swept Field Confocal microscope with 60x oil lens (NA 1.4) and QuantEM model 5125C camera; (ii) for TKO-HEK cells, using an Olympus IX81-DSU Spinning Disk Confocal (SDC) microscope and Andor iXon3 camera (DU-888E-C00-#BV). For GCamP6m-Cytb5 and Bip-mCherry, excitation wavelengths were 488 and 507 nm, and fluorescence emission wavelengths were 507 and 529 nm, respectively. Frames were taken at 1-sec intervals for 5 min. Tm (0.5-2.5 µg/ml, #11089659) was added 20 sec after start of recording.

### Imaging analysis

Ca^2+^ imaging analysis was performed using Image J software program. Bleaching of images taken with SDC microscope was corrected using Exponential Fitting Method in “Bleach Correction” plugin (U.S. National Institutes of Health; http://rsbweb.nih.gov./ij/).

Fluorescence intensity values were plotted as ratios (ΔF/F_0_) of change of fluorescence (ΔF) from region of interest (ROI) (5×5 pixels) divided by mean resting fluorescence (F_0_) prior to Tm addition, vs. recording time. ROIs were defined as active when fluorescence increased ≥2 SD relative to baseline fluorescence.

For co-localisation assays, MOCs were calculated in a 5×5-pixel ROI in the presence or absence of Ca^2+^ release. Average MOC M2 values were plotted vs. average peak Ca^2+^ responses.

### Immunocytochemistry

For immunofluorescence detection, TKO-HEK cells transfected with **p**GCamP6m-Cytb5 were cultured on 12-mm glass coverslips, washed twice with PBS, fixed with 4% paraformaldehyde and 120 mM sucrose in PBS for 15 min at 37°C, permeabilized for 5 min with 0.01% digitonin (#D141) in PBS and sucrose 400 mM, washed 5 min with 1 M KCl, blocked for 45 min in 5% BSA (#A7906) in PBS, incubated overnight at 4°C with anti-S6 ribosomal protein (1:25; #MA5-15123) and Sec61α (1:50; #PA3-014) antibodies, and diluted to indicated concentrations with 5% BSA in PBS. Cells were washed and incubated with Alexa-conjugated secondary antibodies anti-rabbit Alexa Fluor 568 (#A-11011) and anti-mouse Alexa Fluor 405 (#ab175658) for 1 h at room temperature. Images were obtained using Zeiss confocal microscope, model LSM 800.

### Statistical analysis

Statistical analysis was performed using Microsoft Excel V. 14.5.0 and KaleidaGraph V. 4.5.2 software programs. Results are presented as mean ± SEM of 3 or more independent replicates. Significance of differences between means was determined by one-way ANOVA or Tukey’s Multiple Comparison Test in a single step (honestly significant difference test; HSD) as appropriate. Differences with p-values ≤0.05, ≤0.01, and ≤0.001 are indicated respectively by one, two, and three asterisks in the figures.

Graphs were created using Excel 14.5.0, and combined with images using Microsoft PowerPoint 15.5.0.

## Acknowledgements

The authors thank Dr. Exing Wang for invaluable help with imaging, and Dr. Andrea Pellegrini for cell culture technical support.

Calcium images (Figs. 1-4 and Fig. S1) were generated at the Core Optical Imaging Facility, which is supported by UTHSCSA and NIH-NCI P30 CA54174. Images used for Figs. 5-7 and Figs. S2 and S3 were generated at the National Center of Microscopy, Universidad Nacional de Córdoba, Córdoba, Argentina.

This study was supported by grants from: National Institutes of Health (NIH), USA (#RO1AG058778-01A1; Subaward Agreement No. 165148/165147 between UTHSCSA-Instituto Investigación Médica M y M Ferreyra), and Agencia Nacional de Scientific and Technological Promotion, Argentina (ANPCyT, PICT 2017 #0618).

Dr. Constanza Feliziani and Macarena Fernandez were supported by fellowships from The National Scientific and Technical Research Council (CONICET), Argentina.

The wood-whelan research fellowships provided the funds for Dr. Constanza Feliziani’s internship to Dr. James D Lechleiter’s laboratory (Department of Cellular and Structural Biology, University of Texas Health Science Center at San Antonio).

The authors are grateful to Dr. S. Anderson for English editing of the manuscript.

## Author contributions

Conceptualization: M.B., J.D.L. Experiments: C.F., G.Q., D.H, M.F., J.C.P., A.W.P., M.B. Data analysis: C.F., M.B. Manuscript writing: C.F., M.B. Funding acquisition: M.B., J.D.L.

### Competing interests

The authors declare no competing interests.

## Notes

### Competing Interest Statement

The authors have declared no competing interest.

