## Supplemental Information for "Endoplasmic reticulum stress induces a novel Ca^2+^ signalling system initiated by Ca^2+^ microdomains"

### Supplementary information

#### Figure Legends

**Figure S1: Puromycin and anisomycin have opposing regulatory effects on Tm-induced local $Ca^{2+}$ increase.** Human astrocytes were pre-incubated with either puromycin (20 µM, 1 min) or anisomycin (200 µM, 60 min), and added with 0.5 µg/ml Tm. **(a)** Confocal image sequences in pseudo-colour illustrating Tm-induced $Ca^{2+}$ release in control, puromycin-treated, or anisomycin-treated cells. Insets show $Ca^{2+}$ increase events. Scale bar: 30 µm. **(b)** Data for GCamP6-Cytb5 $Ca^{2+}$ increase in terms of $\Delta F/F_0$ were obtained as in Fig. 1 and plotted as a function of time for each condition. Representative data from 3 independent experiments are shown. **(c)** Histograms (mean ± SEM) showing maximal $\Delta F/F_0$ for each condition. *p< 0.05, ***p< 0.0001 (ANOVA, Tukey's HSD test).

**Figure S2: Puromycin regulates Tm-induced local $Ca^{2+}$ increase in TKO-HEK cells.** TKO-HEK cells were added with Tm (2.5 µg/ml) or puromycin (20 µM). Confocal image sequences in pseudo-colour indicate $Ca^{2+}$ release resulting from these treatments. Scale bar: 15 µm (regular image) or 5 µm (magnified image).

**Figure S3:** TKO-HEK cells co-overexpressing GCaMP6-Cytb5 and empty mCherry vector were added with 2.5 µg/ml Tm, and $Ca^{2+}$ confocal images were taken. MOCs were calculated for a 5x5-pixel ROI in which was previously obtained GCamP6-Cytb5 $Ca^{2+}$ increase in terms of $\Delta F/F_0$. MOC **M2** was pooled as sets of values <0.5 and >0.5 (low and high co-localisation, respectively), and average. M2 averages were plotted vs. average peak $Ca^{2+}$ responses. Data were pooled from 3 independent experiments. ns: not significant (ANOVA, Tukey's HSD test).
