## Supplementary figures and images for "Endoplasmic reticulum stress induces a novel Ca^2+^ signalling system initiated by Ca^2+^ microdomains"

### Supplemental Figures 1, 2 and 3

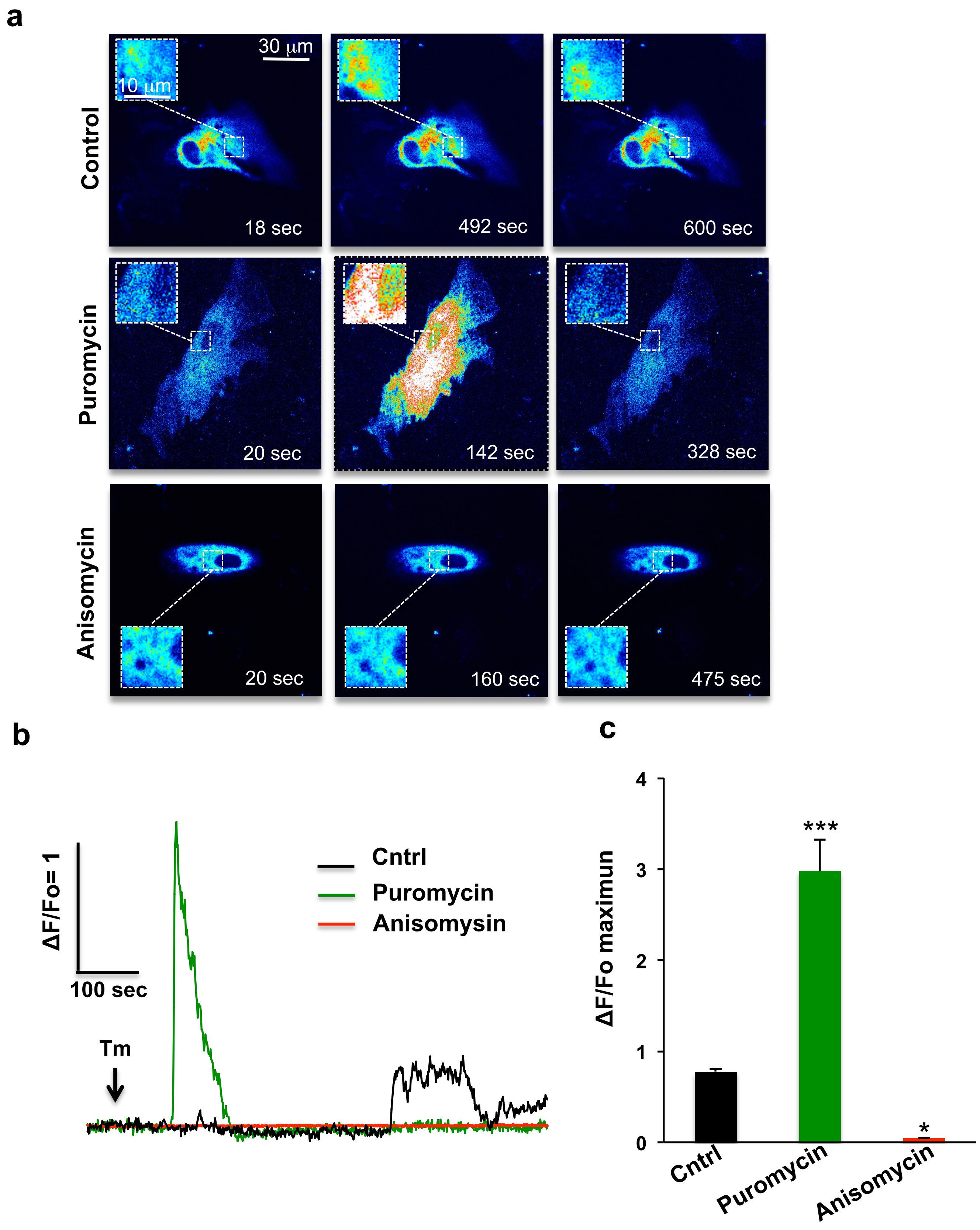

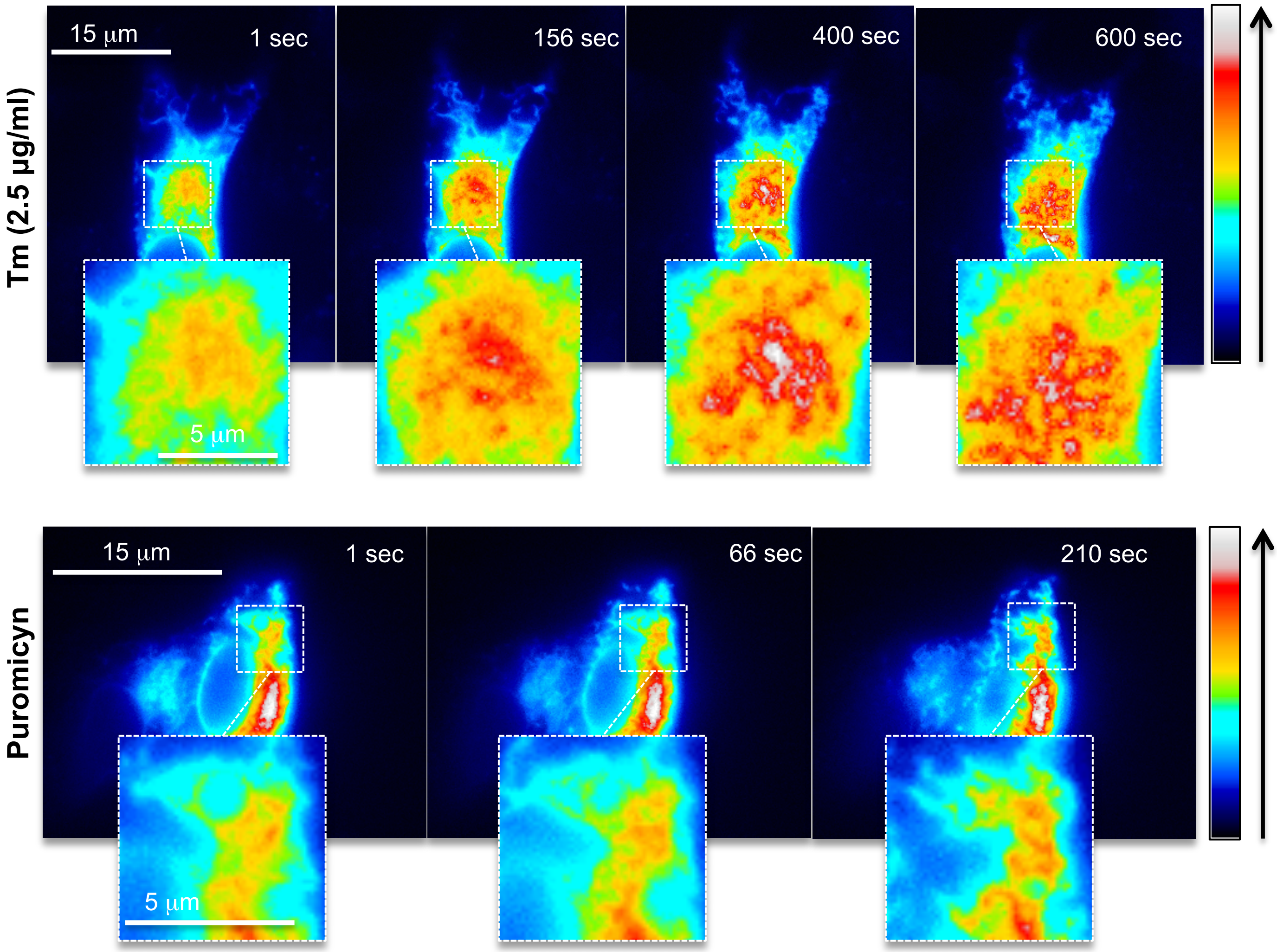

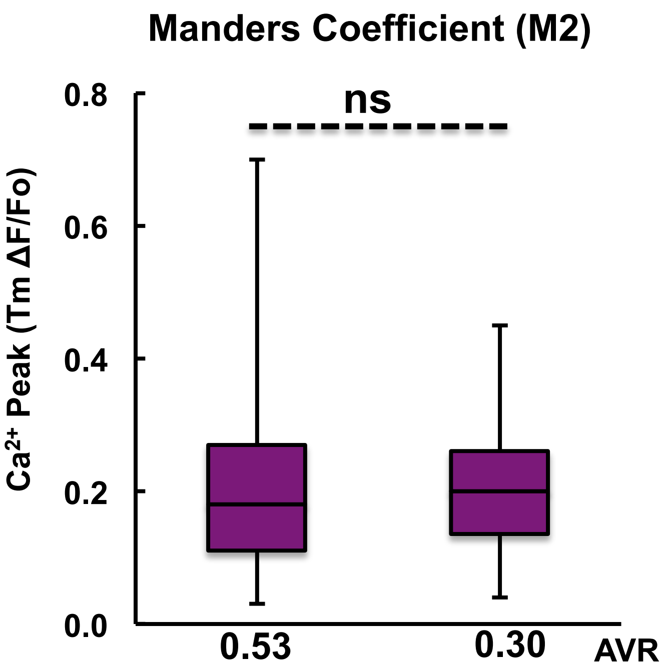
